# Reverse complementary matches simultaneously promote both back-splicing and exon-skipping

**DOI:** 10.1101/2021.02.28.433292

**Authors:** Dong Cao

**Affiliations:** Information Processing Biology Unit, Okinawa Institute of Science and Technology Graduate University, 1919-1 Tancha, Onna, Kunigami, Okinawa, Japan, 904-0495

**Keywords:** circular RNA, back-splicing, exon-skipping, reverse complementary matches, FACS

## Abstract

Circular RNAs (circRNAs) play diverse roles in different biological and physiological environments and are always expressed in a tissue-specific manner. Tissue-specific circRNA expression profile can help understand how circRNAs are regulated. Here, using large-scale neuron isolation from the first larval stage of *Caenorhabditis elegans* (*C. elegans*) followed by whole-transcriptome RNA sequencing, I provide the first neuronal circRNA data in *C. elegans*. I show that circRNAs are highly expressed in the neurons of *C. elegans* and are preferably derived from neuronal genes. More importantly, reverse complementary matches in circRNA-flanking introns are not only required for back-splicing but also promote the skipping of exon(s) to be circularized. Interestingly, one pair of RCM in *zip-2* is highly conserved across five nematode ortholog genes, which show conserved exon-skipping patterns. Finally, through one-by-one mutagenesis of all the splicing sites and branch points required for exon-skipping and back-splicing in the *zip-2* gene, I show that exon-skipping is not absolutely required for back-splicing, neither the other way. Instead, the coupled exon-skipping and back-splicing are happening at the same time.

## Introduction

Circular RNAs (circRNAs) are regulatory RNA molecules that are covalently closed by back-splicing, in which a downstream splice donor is joined with an upstream splice acceptor (Chen, 2020). circRNAs have been reported to bind to microRNAs (the so-called “miRNA sponges”) (Hansen et al., 2013; Memczak et al., 2013) and to act as decoys for proteins to regulate the expression and function of genes (Du et al., 2020; Xia et al., 2018; Zhu et al., 2019). Some circRNAs are translated to functional proteins/peptides through cap-independent mechanisms (Legnini et al., 2017; Pamudurti et al., 2017; Yang et al., 2017; Zhang et al., 2018a; Zhang et al., 2018b; Zheng et al., 2019). circRNAs are always expressed tissue-specifically. Especially, circRNAs are enriched in the brain of several species (Gruner et al., 2016; Rybak-Wolf et al., 2015; Westholm et al., 2014; You et al., 2015), like humans, mice, *Drosophila*, etc. In *C. elegans*, circRNAs are also identified (Ivanov et al., 2015; Memczak et al., 2013) and accumulate during aging (Cortes-Lopez et al., 2018), but such neuronal circRNA profile has not yet been reported.

Back-splicing is a well-regulated process (Chen, 2020). The reverse complementary matches (RCMs) that locate in the flanking introns of circRNA-producing exons promote circRNA formation by bringing the splice sites for back-splicing to proximity. This model was first proposed when a circular transcript was identified in *Sry* in mice, in which a pair of > 15,500 nt RCMs are present in the introns flanking the ∼1,200 nt exon to be circularized (Capel et al., 1993; Gubbay et al., 1992). Subsequent *in vitro* experiments showed that as less as 400 nt of complementary sequences are sufficient enough for the production of *circSry* (Dubin et al., 1995). Genome-wide analysis of RNA-seq data in humans revealed that Alu repeats, which contain RCMs, are enriched in the flanking introns of circRNAs (Jeck et al., 2013). Zhang et al. proved that the orientation-opposite Alu elements and other non-repetitive RCMs sufficiently promote circRNA formation in cultured human cells (Zhang et al., 2014). More importantly, disturbance of RCMs is shown to be an efficient method for circRNA knockout with little effect on cognate linear mRNA (Xia et al., 2018; Zhang et al., 2016). In *C. elegans*, RCMs are abundant in circRNA-flanking introns (Cortes-Lopez et al., 2018; Ivanov et al., 2015), but their roles in circRNA production have not been experimentally tested.

circRNA is found to be correlated to exon-skipping (Barrett et al., 2015; Jeck et al., 2013; Kelly et al., 2015; Surono et al., 1999; Zaphiropoulos, 1996, 1997). In early years, sporadic examples showed that some circRNA-producing genes generate linear transcripts that skip the exons to be circularized (Surono et al., 1999; Zaphiropoulos, 1996, 1997). Later, systematic analysis of RNA-seq data in human cells found a global correlation of exon-skipping with exon circularization (Kelly et al., 2015). In *Schizosaccharomyces pombe*, Barrett et al. showed that circRNA could be produced from an exon-containing lariat intermediate produced by exon-skipping (Barrett et al., 2015). Given that the correlated exon-skipping and circRNA formation use the same pair of introns, it is possible that RCMs in circRNA-flanking introns also regulate exon-skipping.

Here, using *C. elegans* as the model organism, I provide the first circRNA profile in its neurons by large-scale neuron isolation followed by whole-transcriptome RNA-seq. Hundreds of novel circRNAs were annotated, mainly from the sorted neuron samples. As in other organisms, circRNAs are also abundant in the neurons of *C. elegans*. Dozens of neuron-enriched and neuron-depleted circRNAs are identified. Interestingly, circRNA levels show strong correlations to their cognate linear mRNA levels between the sorted neurons and whole worm samples. By deleting one of the RCMs in multiple circRNA genes, I experimentally validated that RCMs are required for circRNA formation in *C. elegans*. Moreover, RCMs also promote correlated exon-skipping. Notably, one pair of RCM in the *zip-2* gene is highly conserved across five nematode species and likely has conserved roles in promoting exon-skipping. Finally, through *in vivo* one-by-one mutagenesis of splicing sites and branch points of exon-skipping and back-splicing, I show that RCM-promoted back-splicing and RCM-promoted exon-skipping are happening together to produce the circular and the skipped transcripts.

## Results

### 1. Successful neuron isolation for circRNA detection by RNA-seq

Currently, there are no available neuronal circRNA data in *C. elegans*, mainly due to challenges in obtaining enough neuron samples from the tiny worms which have no obviously compartmentalized “brain” tissue. The most common method to obtain neuron cells from *C. elegans* is by the “labeling-dissociation-sorting” method (Figure 1A) (Christensen et al., 2002; Deffit et al., 2017; Fox et al., 2005; Kaletsky et al., 2016; Spencer et al., 2014; Zhang et al., 2011; Zhang et al., 2002), in which target neurons are labeled by fluorescent protein and, after mild dissociation of the worms, labeled neurons are collected by fluorescence-activated cell sorting (FACS). This method can obtain target neurons in high purity and is used to detect gene expression in single neurons to all the neurons. However, due to the low efficiency of the dissociation (Spencer et al., 2014; Zhang et al., 2011), total RNA obtained from sorted cells is limited. Hence this method is only used for mRNA detection, either by microarray or RNA-seq (Deffit et al., 2017; Kaletsky et al., 2016; Spencer et al., 2014; Von Stetina et al., 2007; Watson et al., 2008). By optimization of previous protocols (Zhang et al., 2011), I aim to improve the final total RNA yield to hundreds of nanograms for circRNA detection by whole-transcriptome RNA-seq.

**Figure 1.**
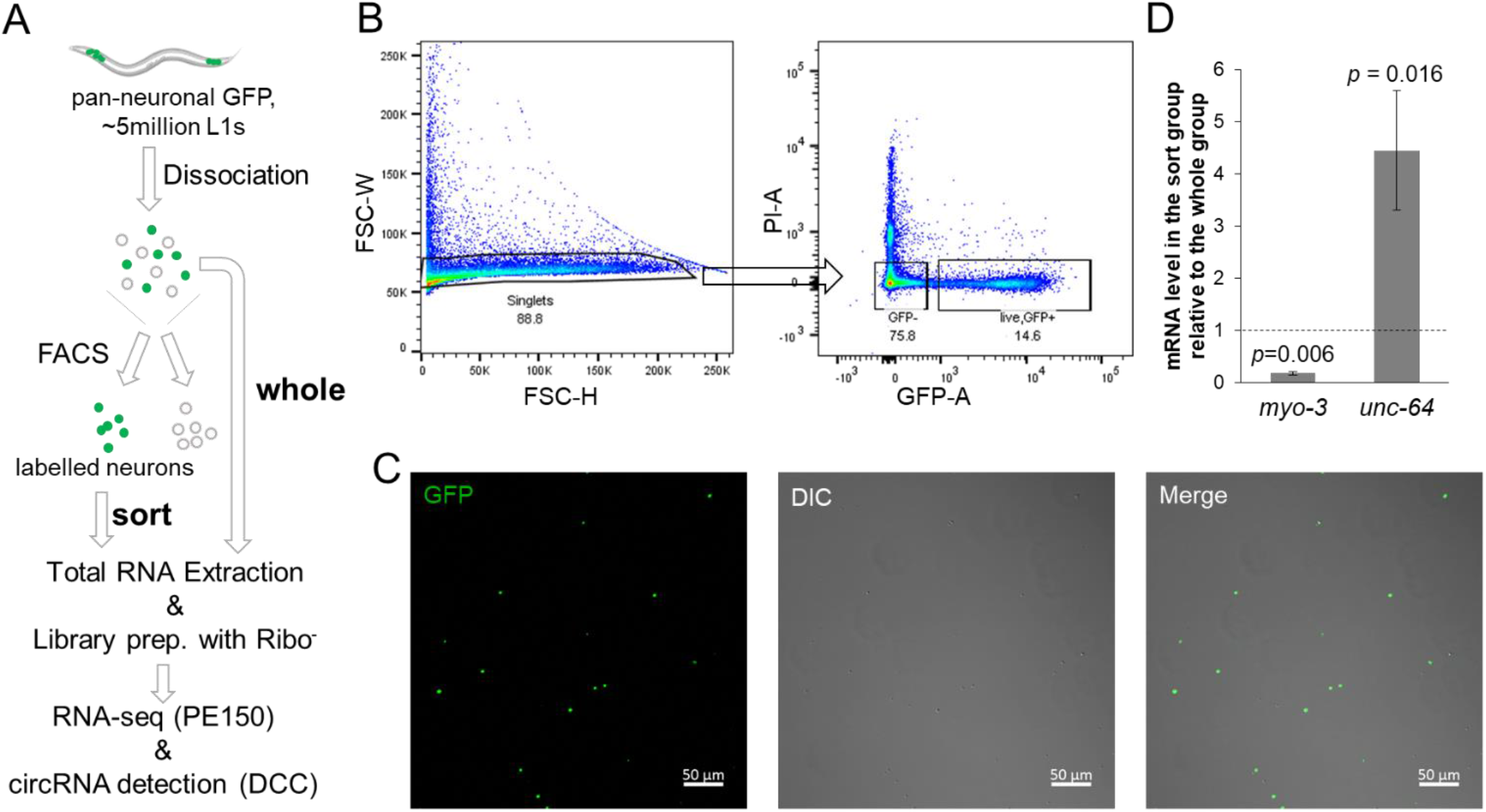
Large scale neuron isolation from *C. elegans* for circRNA detection. (A) Workflow of neuron isolation and circRNA detection by RNA-seq. (B) Gating strategy for FACS: Forward scatter width (FSC-W) was plotted against forward scatter height (FSC-H) to select singlet cells (88.8%), which were then used for the selection of GFP-positive and PI-negative cells (14.6%) for sorting (C) Confocal images of sorted GFP-positive neurons. Scale bar: 50 μm. (D) ddPCR results showing the relative levels of two genes (*myo-3* and *unc-64*) in the sort group compared with those in the whole group. Error bars stand for standard deviations of three biological replicates. *P* values are ratio paired *t*-test.

Here, using a strain (NW1229) with pan-neuronal green fluorescent protein (GFP) expression, I found that by shortening the time of SDS-DTT treatment (from 2 min to 1.5 min) and washing (from 5 x 1.0 min to 5 x 40 sec) as well as increasing the time of mechanical disruption (from 10 min to 15 min) (Figure S1A), cell yield could be improved. After dissociation, the cell suspension was stained with propidium iodide (PI) to label dead/damaged cells and then subjected to FACS. GFP positive singlet cells were sorted (Figure 1B and Figure S1B). The majority of sorted cells showed GFP fluorescence when observed under a confocal microscope (Figure 1C). Consistent with previous findings, neurites can grow out after culture of sorted neuron cells (Figure S1C) (Spencer et al., 2014; Zhang et al., 2011). To further confirm the effectiveness of sorting, the levels of two marker genes (*myo-3* and *unc-64*) were quantified by digital droplet PCR (ddPCR). As expected, the neural syntaxin *unc-64* was highly enriched in the sorted cells, whereas the muscle gene *myo-3* was depleted (Figure 1D).

Using this optimized protocol (see “Materials and methods”), 200 - 500 ng total RNA was obtained from cells sorted from ∼ 1.5 - 5 million L1 worms (the sort group). RNA samples from dissociated worms before sorting were also prepared for comparison (the whole group, Figure 1A). For RNA-seq, 150 ng total RNA from three independent trials of the sort group and the whole group was used as input for library preparation with ribosomal RNA removal and first-strand cDNA synthesis using random hexamers. More than 45 million 150 nt paired-end reads were obtained for each sample. Differentially expressed genes between the two groups were analyzed by DESeq2 (Love et al., 2014). Consistent with the ddPCR results (Figure 1D), *myo-3* was significantly depleted, while *unc-64* was significantly enriched in the sort group compared with the whole group (Figure S1D). The significantly upregulated genes (Table S4) in the sort group were searched in WormExp (Yang et al., 2016) (https://wormexp.zoologie.uni-kiel.de/wormexp/) to identify whether these genes overlap with previous results of neuronal genes. As expected, the resulted top three datasets were all pan-neural enriched genes determined by microarray analysis of sorted neurons (Figure S1E) (Von Stetina et al., 2007; Watson et al., 2008), indicating the RNA-seq results from sorted samples successfully revealed the gene expression pattern in the neurons.

### 2. circRNAs are highly expressed in the neurons

circRNA annotation from the RNA-seq data was carried out using the DCC pipeline (Cheng et al., 2016). Prior to filtering, 6475 circRNAs were identified with at least one junction read across six samples, with 4786 novel circRNAs when compared with circRNAs of *C. elegans* in two databases: circBase (Glazar et al., 2014) and CIRCpedia v2 (Dong et al., 2018) (Figure S2A). The results were filtered with a cutoff of at least three junction reads in each group, which gave 1452 circRNAs derived from 1010 genes and 29 not-annotated loci (Figure 2A, Table S5). Of the filtered circRNAs, the majority of the identified junction reads were from exon-to-exon joining (Figure S2B). The filtered circRNAs were compared with a published dataset of circRNAs in aging worms (Cortes-Lopez et al., 2018), which showed 450 overlapped circRNA (Figure S2C). The novel circRNAs identified in my dataset were mainly from the sorted group, suggesting that sequencing from sorted neuron samples was helpful to identify circRNAs that may not be detected using whole-worm samples. Gene ontology (GO) enrichment analysis of the circRNA-producing genes showed that terms related to the neuronal functions were significantly enriched, including neurogenesis, synaptic signaling, etc. (Figure 2B).

**Figure 2.**
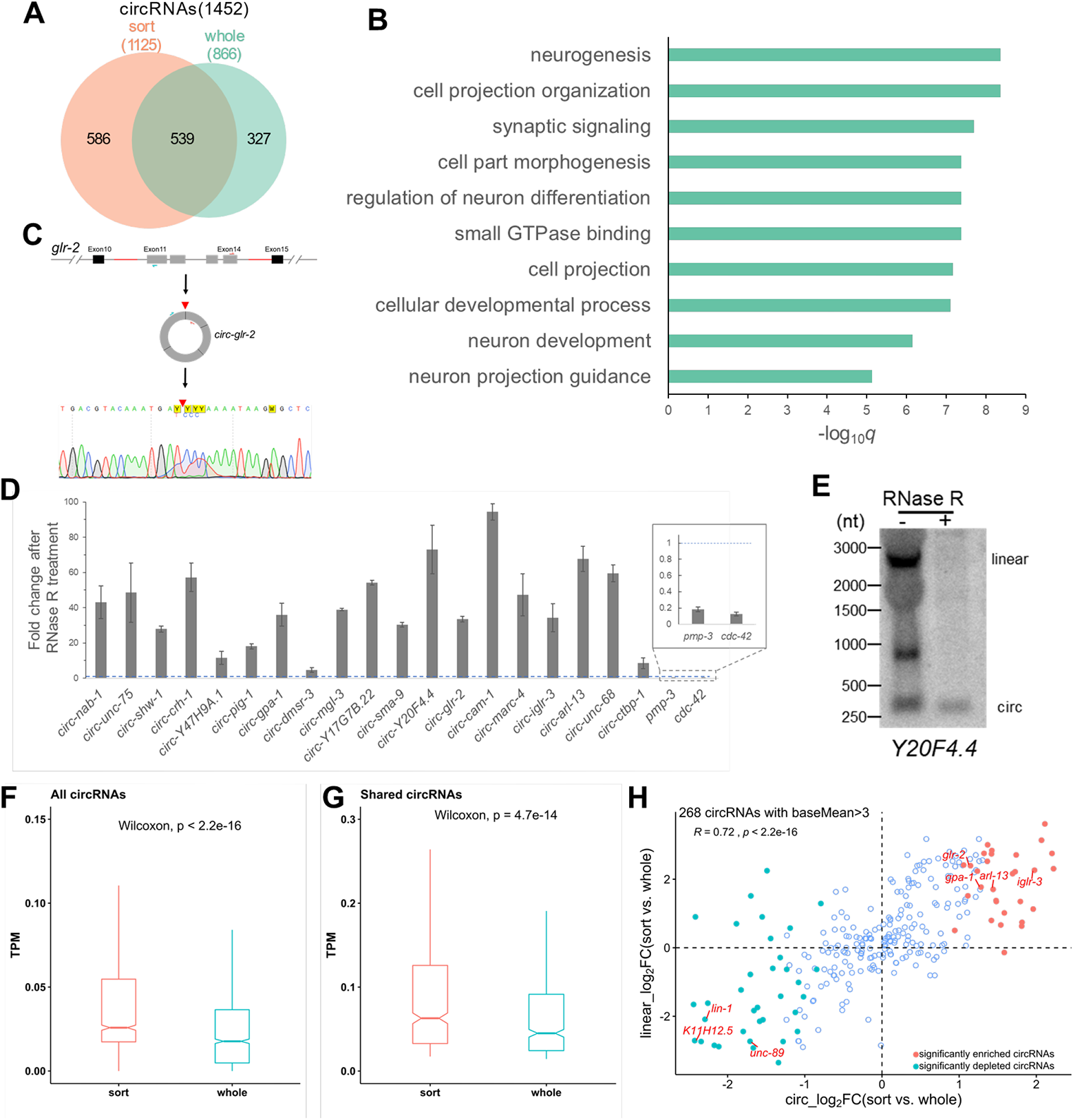
circRNAs are enriched in the neurons of *C. elegans*. (A) Overlap of circRNAs detected in the “sort group” and the “whole group”. (B) Top 10 enriched gene ontology (GO) terms of circRNA-producing genes. (C) Scheme showing amplification of back-splicing junction of a circRNA from *glr-2* using divergent primers. Amplified sequences are confirmed by Sanger sequencing. The red triangle denotes the joint site. (D) RT-qPCR results of the fold changes of circRNAs and two linear mRNAs (*pmp-3* and *cdc-42*, inset) after RNase R treatment. The blue dashed line shows one-fold change. Error bars are the standard deviations of three biological replicates. (E) Northern blot detection of *Y20F4*.*4* transcripts in total RNA (20 μg) without or with RNAse R treatment, using probes that hybridize to both linear and circular transcripts. (F, G) TPM (transcripts per million reads) comparison of all circRNAs (F) and shared circRNAs (G) between the “sort group” and the “whole group”; *p* values are paired Wilcoxon test. (H) Scatter plot showing the fold changes of 268 circRNAs with baseMean > 3 versus fold changes of their corresponding linear mRNAs. The Pearson correlation coefficient (*R*) and *p* value (*p*) are shown. Significantly differentially expressed circRNAs are shown by colored dots. Names of several circRNA genes are labeled.

Two strategies were used to validate the annotated circRNAs by DCC: 1) Amplification of the back-spliced junction sequences by RT-PCR using divergent primers followed by Sanger-sequencing (Figure 2C). Eighteen out of 19 selected circRNAs, including seven novel circRNAs, were confirmed with the back-splicing junction (BSJ) sequences (Figure S2D). 2) Enrichment quantification by RT-qPCR after RNase R treatment. Since there are no ends in circRNAs, they often show resistance to degradation after treatment with RNase R. As expected, while two linear mRNAs (*pmp-3* and *cdc-42*) were depleted after RNase R treatment, all the circRNAs were enriched (Figure 2D). The resistance to RNase R was also confirmed by northern blot, which showed that while linear transcript was not detected after RNase R treatment, circRNA from *Y20F4*.*4* was still detected (Figure 2E). Together, these results show that circRNA annotation is of high accuracy.

Of the 1452 circRNAs, more circRNAs (1125/1452) were found in the sort group, with 539 identified in both groups and 586 only in the sort group (Figure 2A). Next, the abundances of circRNAs in the sort group and the whole group were compared to check whether circRNAs were highly expressed in the neurons of *C. elegans* or not. TPM (transcripts per million reads) values were used for comparison. The principal component analysis (PCA) plot of circRNA TPM showed a clear separation between the two groups (Figure S2E), suggesting different circRNA profiles between them. For all the circRNAs in both groups, circRNAs in the sort group showed significantly higher TPM values than in the whole group (Figure 2F, *p* < 2.2e-16, paired Wilcoxon test), indicating circRNAs were enriched in the sort group. The same trend was also observed for the shared 539 circRNAs in both groups (Figure 2G, *p* = 4.7e-14, paired Wilcoxon test).

Next, differentially expressed circRNAs between the sort and the whole group were analyzed, trying to identify neuron-enriched circRNAs. Using BSJ read numbers as input for DESeq2 and adjusted *p* value < 0.05, 31 circRNAs were found significantly enriched, and 35 circRNAs were significantly depleted in the sort group (Figure S2F, Table S6). I asked whether these circRNAs were also derived from neuronal genes or not. The fold changes of circRNAs between the sort and the whole group were plotted against the fold changes of their cognate linear mRNAs. Here, a cutoff of baseMean (given by DESeq2) bigger than 3 was used, which contained 268 circRNAs, including all the significantly differentially expressed circRNAs (Figure 2H). The results showed a strong positive correlation (Figure 2H, Pearson’s correlation coefficient *R* = 0.72, *p* < 2.2e-16), which suggested that at the L1 stage of *C. elegans*, neuronal circRNAs were more likely to be derived from neuronal genes. When all circRNAs were considered, they still showed a moderately strong positive correlation (Figure S2G, Pearson’s correlation coefficient *R* = 0.51, *p* < 2.2e-16).

In summary, the first neuronal circRNA profiles in *C. elegans* have been obtained, which show that circRNAs are highly expressed in the neurons and positively correlated with their cognate mRNAs.

### 3. RCMs are required for circRNA production

Next, features of circRNA-flanking introns were analyzed. Similar to previous findings (Cortes-Lopez et al., 2018; Ivanov et al., 2015), introns that flank circRNA-producing exons were much longer than average, and much more RCMs were identified when compared with flanking introns of non-circular exon controls (exon 2 and exon 8) (Figure 3A and 3B). Lengths of the best-matched RCMs in circRNA introns were also much longer than those in introns flanking control exons (Figure S3A). Although a previous study showed that RCMs could predict the existence of circRNAs (Ivanov et al., 2015), the role of RCMs in circRNA formation has not been experimentally confirmed in *C. elegans*. Here, six circRNA genes with RCMs in flanking introns were chosen, and one RCM in each gene was deleted using CRISPR-Cas9 (See materials and methods, Figure 3C and Figure S3B). Which RCM was selected for deletion depends on its position in that intron and the existence of high-specificity guide RNA (gRNA) sites around RCM sequences. For example, in *glr-2*, the downstream RCM is very close to the 3’ splice site, so the RCM in the upstream was chosen for deletion (Figure S3D). The gRNA sequences and recombinant single-strand oligo DNAs used for RCM deletions are listed in Table S3. As expected, all the circRNAs were either undetectable or reduced to a very low level after the removal of one RCM in the flanking introns (Figure 3D and Figure S3C), proving that RCMs in *C. elegans* strongly promote circRNA formation. Of note, in some of the chosen genes, the linear mRNA levels were altered in RCM deletion mutants.

**Figure 3.**
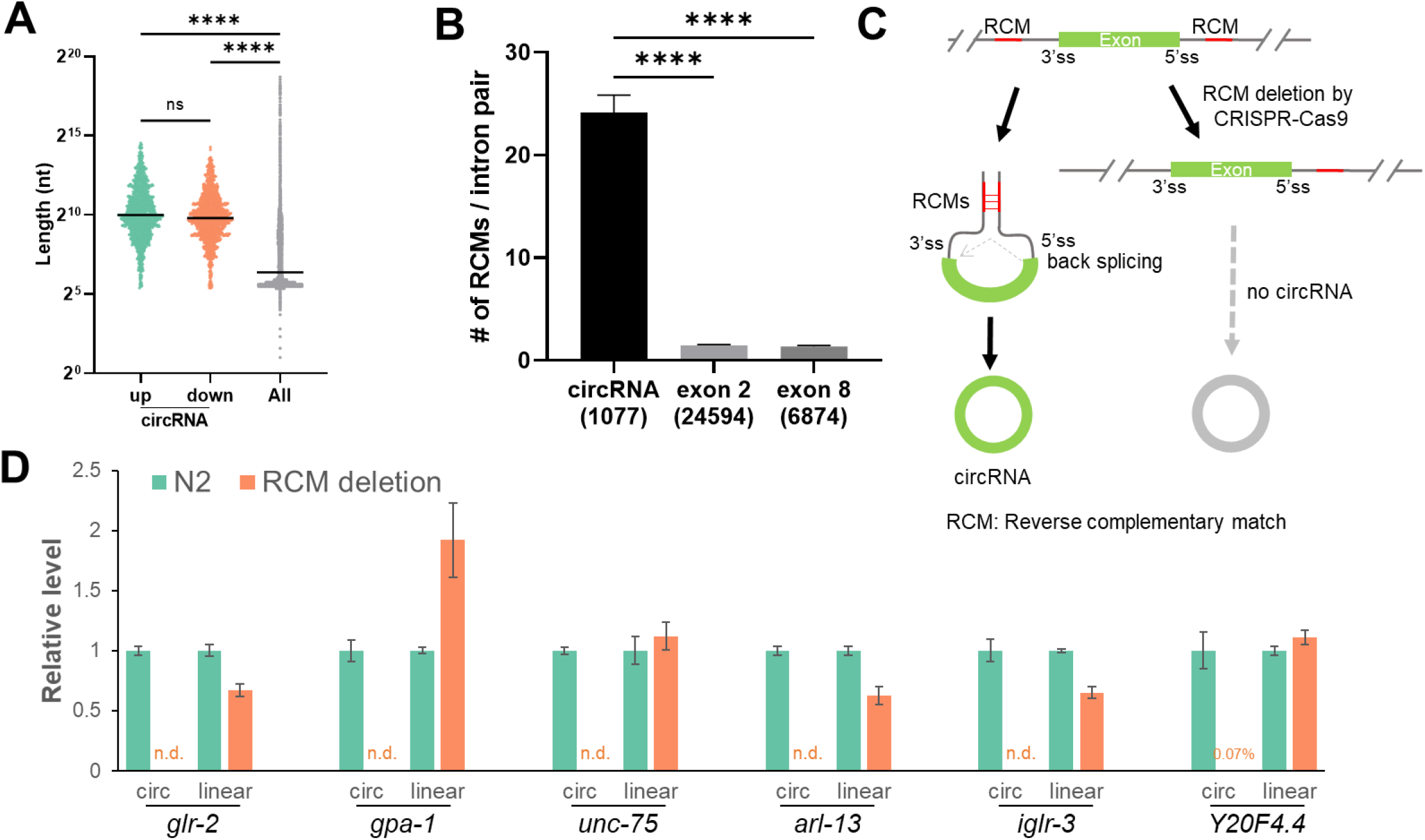
RCMs are required for circRNA production. (A) Length distributions of introns flanking circRNA-producing exon(s), compared with the lengths of all introns. (B) Average number of RCMs in one pair of flanking introns of circRNA compared with those in non-circRNA control exons (exon 2 and exon 8). Values are shown as mean ± SEM. Numbers in the brackets are numbers of intron pairs used for analysis. *p* values in A & B are from Kruskal-Wallis test with Dunn’s post-hoc test for multiple comparisons. ****, *p* < 0.0001. (C) Schematic plot showing that RCMs promote circRNA production and the strategy to disturb one of the RCMs by CRISPR-Cas9. (D) Quantification of linear mRNA and circRNA in wildtype N2 strain and RCM deletion mutant strains of six circRNA genes. Error bars are the standard deviations of three biological replicates. n.d.: not detected (Ct values not determined or bigger than those in no-template controls (NTC)).

### 4. RCMs promote both back-splicing and exon-skipping

circRNA production has been correlated with exon-skipping that skips the circularized exon(s) (Kelly et al., 2015; Surono et al., 1999; Zaphiropoulos, 1996, 1997). In the RNA-seq data of this study, reads mapped to the skipped junctions can be identified in some circRNA genes (Figure S4A and S4B), suggesting the existence of skipped transcripts. For *zip-2* and *Y20F4*.*4*, RT-PCR using primers that bracket circRNA-producing exon(s) gave two bands, of which the longer ones were full-length transcripts and the shorter ones were confirmed to be the skipped transcripts (Figure 4A and Figure S4E). For some other circRNA genes, the skipped transcripts could be amplified in a two-round PCR, in which the corresponding skipped transcripts were gel-cut purified after first-round PCR, which were used as templates for a second-round PCR (Figure S4C and S4D). In total, skipped transcripts were confirmed in 6 out of 7 chosen circRNA genes (Figure S4E).

**Figure 4.**
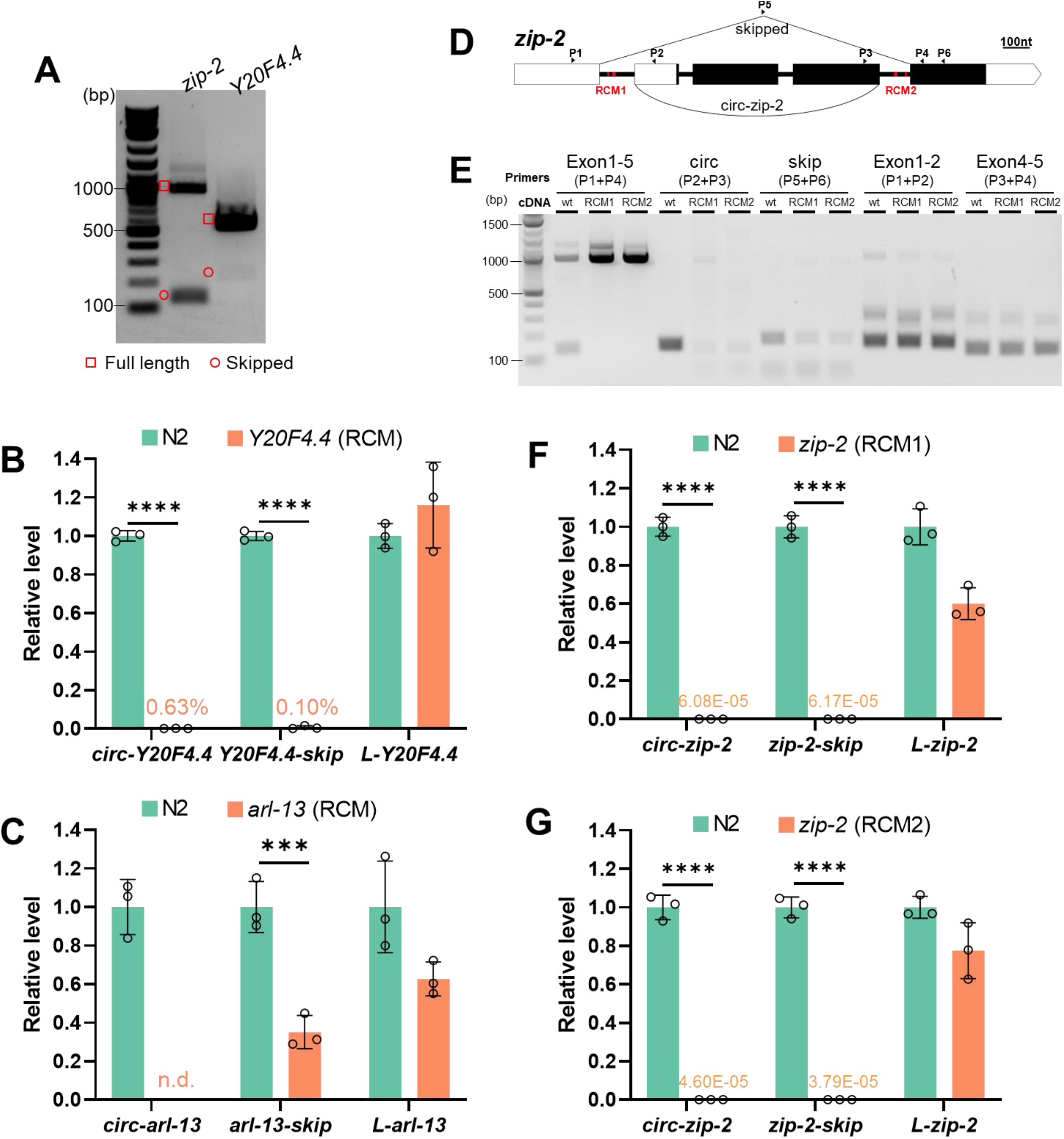
RCMs promote both back-splicing and exon-skipping. (A) RT-PCR detection of full-length transcripts and skipped transcripts in *zip-2* and *Y20F4*.*4*. (B) RT-qPCR quantification of *Y20F4*.*4* transcripts in wildtype and RCM-deleted *Y20F4*.*4* strain. (C) RT-qPCR quantification of *arl-13* transcripts in wildtype and RCM-deleted *arl-13* strain. (D) Illustration of the gene structure of *zip-2*. P1-P6: positions of primers. Black rectangles indicate coding regions, and white parts are untranslated regions (UTRs). RCM areas are in red. (E) RT-PCR detection of transcripts from *zip-2* gene in wildtype N2 strain and RCM-deleted strains. (F, G) RT-qPCR quantification of *zip-2* transcripts (circular: *circ-zip-2*, skipped: *zip-2-skip*, full length linear: *L-zip-2*) in RCM-deleted strains compared with wildtype N2 strain. (B,C F & G) Results are normalized to levels in N2 strain using *pmp-3* as the reference gene. Error bars are the standard deviations of three biological replicates. ***, *p* < 0.001, ****, *p* < 0.0001, two-tail Student’s *t-*test.

Previous studies have shown that conserved complementary sequences in introns are associated with exon-skipping (Miriami et al., 2003). Complementary sequences in different introns regulate mutually exclusive splicing (Graveley, 2005; May et al., 2011; Yang et al., 2011). Then whether RCMs are also required for exon skipping is checked. In *Y20F4*.*4*, the skipped transcript was strongly reduced after removing the upstream RCM (Figure 4B and Figure S5A). In *arl-13*, the skipped transcript was downregulated in the downstream RCM-deleted mutant (Figure 4C and Figure S5B). In *zip-2*, two pairs of perfectly matched RCMs, 7 nt and 13 nt in length, respectively, were identified (Figure 4D and Figure S5C). Deletions of the RCMs in intron 1 or intron 4 were achieved by CRISPR-Cas9 (Figure S5C). Canonical splicing of intron 1 and intron 4 were not affected by RCM deletions (Figure 4E, Exon 1-2 & Exon 4-5). However, although the circRNA and skipped transcript can be detected in the RCM-deleted strains, their production seemed not as efficient as in wildtype strain (Figure 4E, circ & skip). Quantification of the levels of the 3 transcripts of *zip-2* (circular, skipped, full-length linear) showed that while full-length linear *zip-2* was only slightly affected, the production of both the circRNA and the skipped transcript was dramatically reduced in both RCM-deleted mutant strains (Figure 4F and 4G). Together, these findings suggest that RCMs in the flanking introns of circRNA-producing exon(s) also promote the skipping of these exon(s).

### 5. RCM sequences in *zip-2* are highly conserved across several nematode species

Previous studies suggest that competing RNA secondary structures formed by base-pairing between introns that regulate mutually exclusive splicing are evolutionally conserved (Hong et al., 2020; Yue et al., 2017). I then checked whether RCM sequences in *zip-2* are conserved or not. Ortholog genes of *zip-2* exist in five nematode species (*C. elegans, C. brenneri, C. briggsae, C. japonica*, and *C. remanei*). These *zip-2* genes have similar gene structures (Figure S6). Sequences in the upstream introns and downstream introns of these *zip-2* genes were aligned. Of the two pairs of RCMs in *zip-2* of *C. elegans*, the 13-nt RCMs are highly conserved across the five nematode species, both in the upstream introns and the downstream introns (Figure 5A and 5B). Using available splicing data on WormBase, transcripts that skip exons bracketed by the conserved RCMs were found in all these *zip-2* genes (Figure S6), suggesting the conserved RCMs possibly promote the conserved exon-skipping in all these *zip-2* genes.

**Figure 5.**
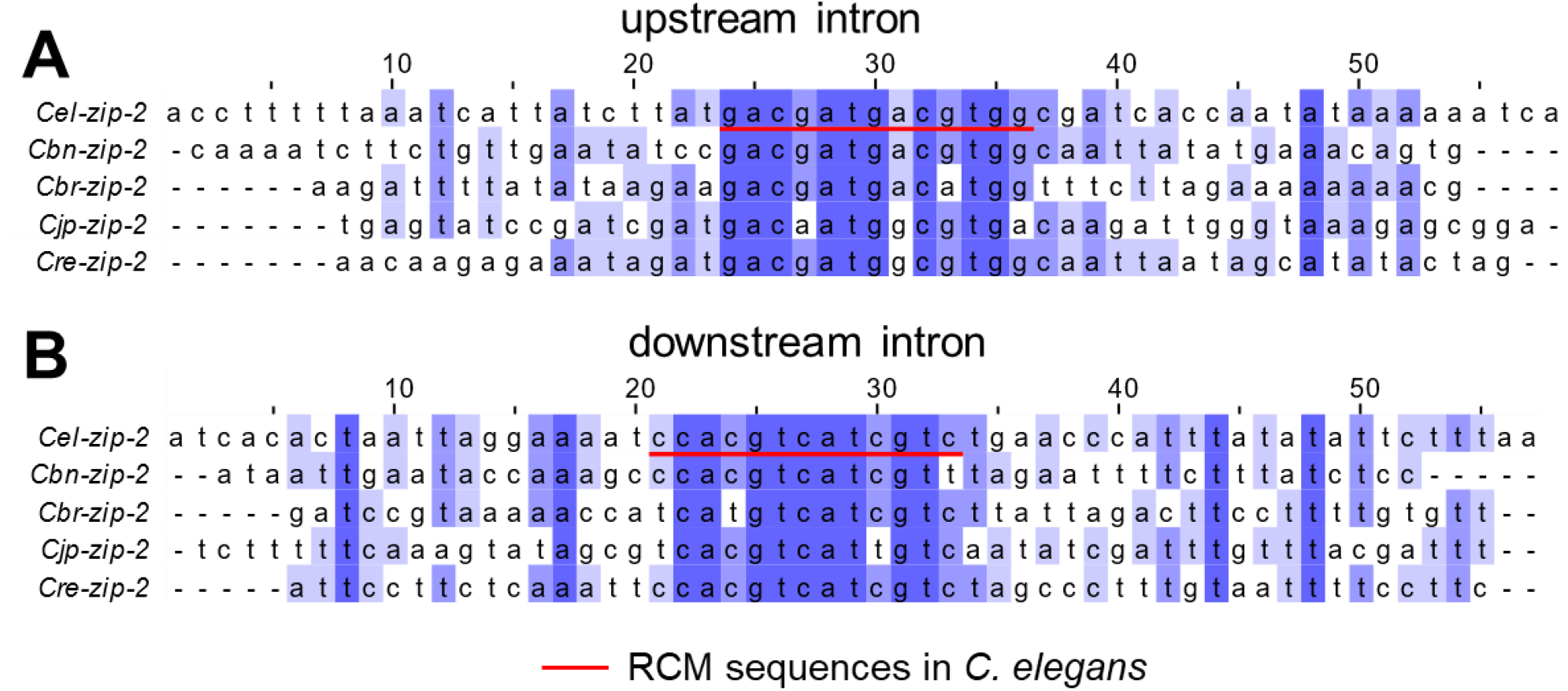
RCM sequences in *zip-2* are highly conserved across several nematode species. (A, B) Alignment of intronic sequences in ortholog *zip-2* genes in indicated nematode species. Red lines underline RCM sequences in *zip-2* of *C. elegans*.

### 6. RCMs do not promote exon-skipping through back-splicing, neither the other way

Current knowledge suggests that RCMs promote circRNA formation by bringing the splicing sites for back-splicing in close proximity. Since the correlated back-splicing and exon-skipping use the same pair of introns, it is possible that RCMs also bring the splice sites for exon-skipping together. In principle, the y-shaped intermediate of back-splicing could be further spliced to form the corresponding skipped transcripts. Moreover, a previous study has shown that circRNA can be produced through a lariat intermediate produced by exon-skipping (Barrett et al., 2015). Whether RCMs promote exon-skipping first or back-splicing first? There are three possibilities: 1). RCMs promote back-splicing first; 2). RCMs promote exon-skipping first; 3). RCMs promote both back-splicing and exon-skipping at the same time (Figure 7). In order to clarify the three possibilities, the 4 splice sites (ss) and 2 branch points (BP) in intron 1 and intron 4 of *zip-2* were mutated one by one. The 5’ss in intron 1, BP, and 3’ss in intron 4 are used for exon-skipping; hence these sites are named skip-5’ss, skip-BP, and skip-3’ss, respectively. Similarly, BP and 3’ss in intron 1 and 5’ss in intron 4 were named circ-BP, circ-3’ss, and circ-5’ss, respectively (Figure 6A). For ss mutation, the conserved AG or GT nucleotides were deleted, and some possible cryptic splice sites nearby were mutated (Figure S7A and S7B). For BP mutation, since there is little information about BP sites in *C. elegans* (Zahler, 2012), all A nucleotides were changed to G nucleotides in the 3’ half of intron 1 and intron 4, without disturbing the RCM sequences. (Figure S7A and S7B). The results showed that mutation of ss and BP for exon-skipping sufficiently abolished *zip-2-skip*. However, *circ-zip-2* was still produced in these mutant strains (Figure 6B). For mutations of ss/BP required for back-splicing, circ-3’ss mutation produced a circRNA using a noncanonical AA site (Aroian et al., 1993) (Figure 6B, Figure S7B, and S7C). circ-5’ss and circ-BP mutation both blocked circRNA formation, but the skipped product can still be detected (Figure 5B). These results suggest that in *zip-2*, exon-skipping is not absolutely required for back-splicing and *vice versa*. RCMs can promote both exon-skipping and back-splicing directly at the same time.

**Figure 6.**
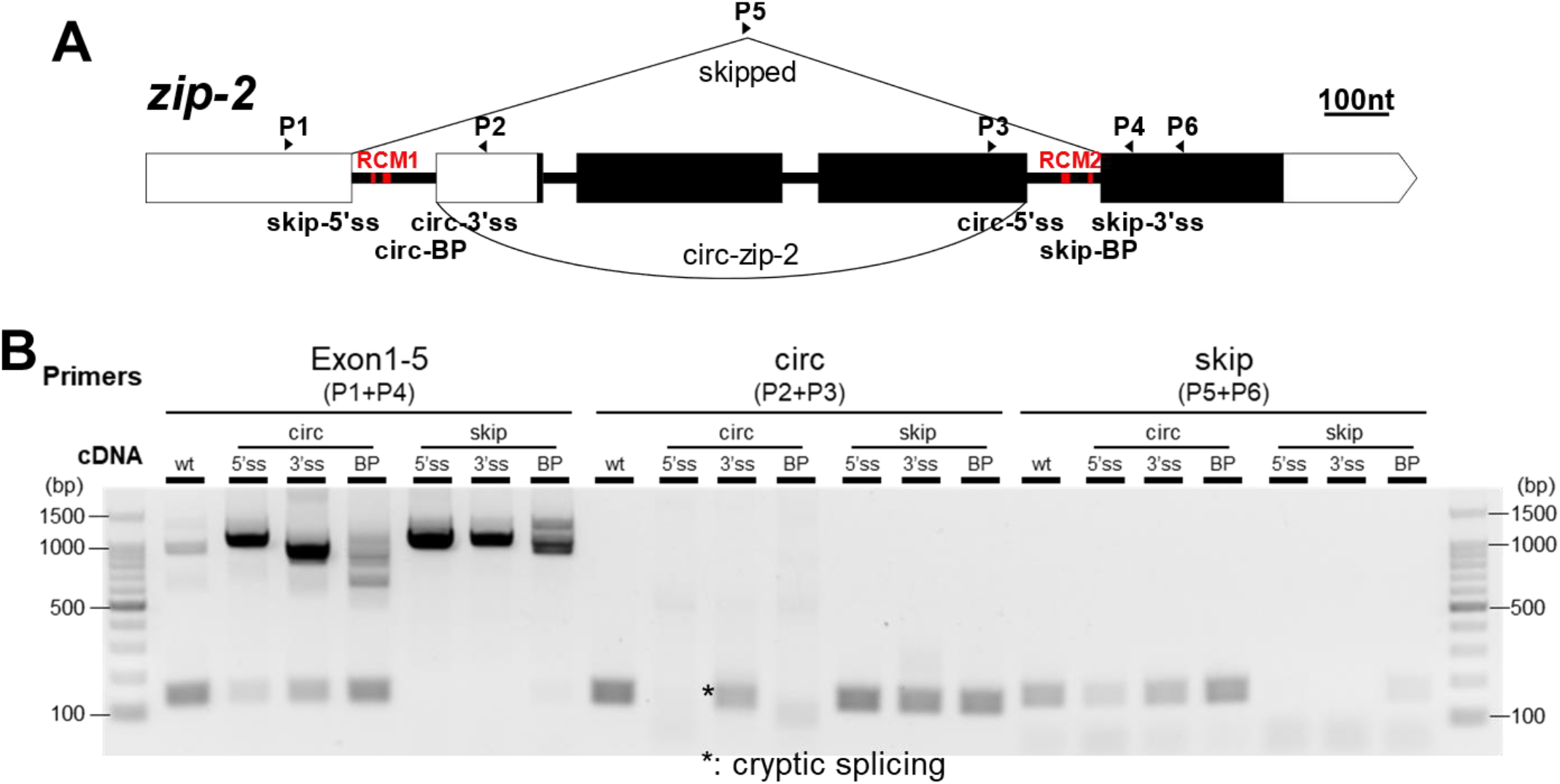
RCMs do not promote exon-skipping through back-splicing, neither the other way. (A) Gene structure of *zip-2*. P1-P6: positions of primers. Positions of splicing sites and branch points that are required for back-splicing and exon-skipping are labeled. Positions of RCMs are in red. (B) RT-PCR detection of *zip-2* transcripts in wildtype N2 strain and strains with mutated ss or BP. Note the cryptic splicing in circ-3’ss strain.

**Figure 7.**
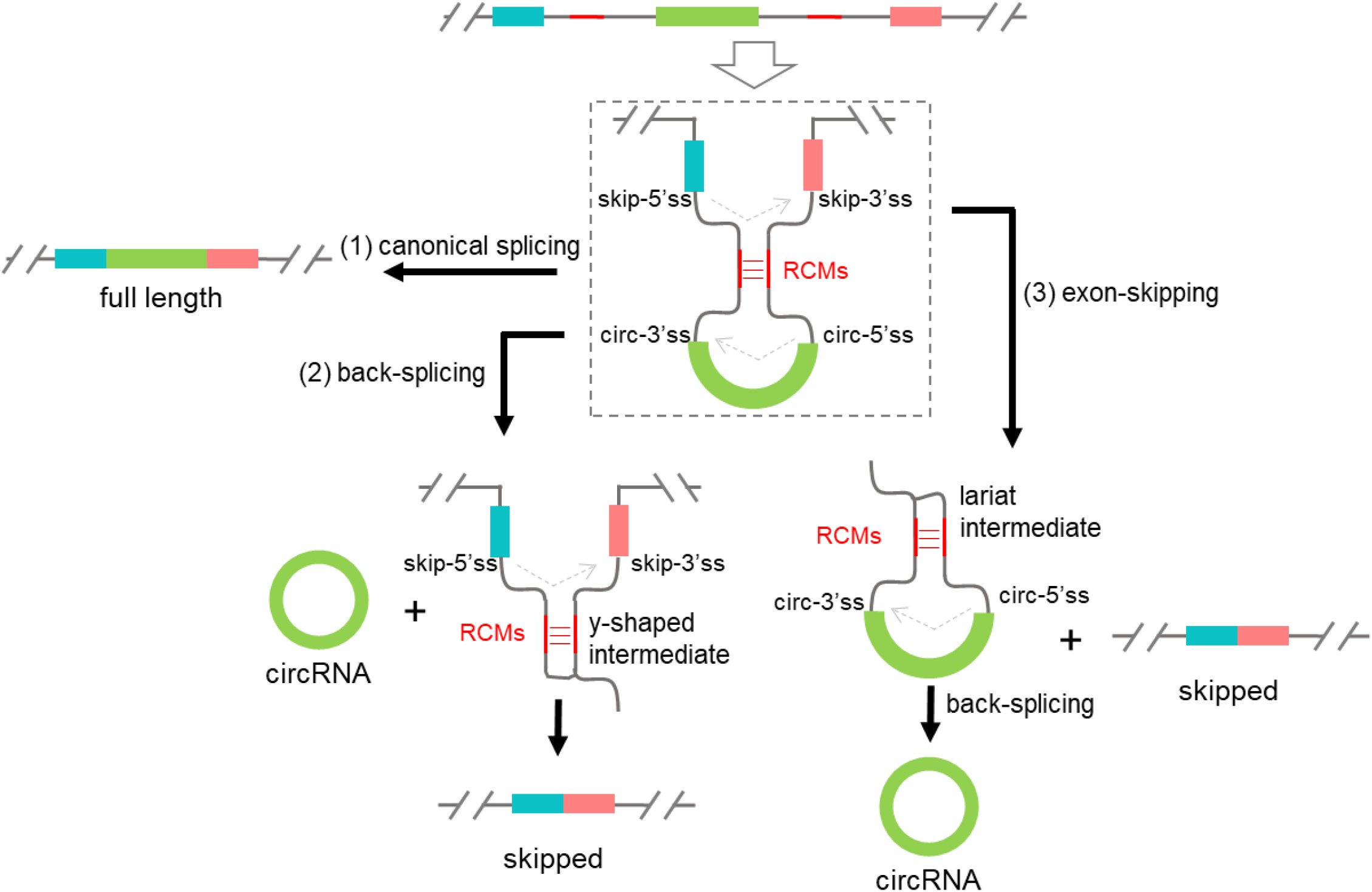
A proposed model that RCMs promote both back-splicing and exon-skipping at the same time. (1) Canonical splicing to form full-length linear mRNA. (2) RCMs facilitate circRNA formation by bringing splice sites for back-splicing sites together. The y-shaped intermediate is further spliced to form the skipped transcript. (3) RCMs promote exon-skipping by bringing splice sites for exon-skipping together. The lariat intermediate is further backspliced to form circRNA. RCMs in the y-shaped intermediate and the lariat intermediate may help the second splicing steps. The three pathways compete with each other to regulate the levels of the three types of transcripts.

## Discussion

In this study, I optimized a method for large-scale neuron isolation from L1 worms. The amount of obtained RNA from sorted neurons was increased for the first time to hundreds of nanogram scale, making the detection of circRNA by whole-transcriptome RNA-seq more reliable. Using this method, I provided the first neuronal circRNA profiles in *C. elegans* and found that circRNAs were abundant in the neurons. Interestingly, circRNAs showing higher levels in the neurons tend to be derived from genes that also show higher expression in the neurons (Figure 2H). The time between egg to L1 is the first main period of neuron development, and at the time of hatching, the majority (222/302) of neurons are already formed (Altun and Hall, 2011). The high levels of these circRNAs may be due to the active expression of their parental genes for neuron development at the L1 stage.

RCMs are abundant in circRNA introns of *C. elegans*. For the first time, I validated that RCMs are required for circRNA formation in multiple circRNA genes. This provides a good method to knockout (KO) circRNAs in *C*. elegans. Especially, RNA interference (RNAi) in *C. elegans* produces secondary short interfering RNAs (siRNAs) that recognize sequences other than the primary targets in the same genes (Billi et al., 2014), which probably makes circRNA-specific knockdown (KD) by RNAi not working in *C. elegans*. Except for developing the Cas13-based KD method (Li et al., 2020) in *C. elegans*, disrupting RCMs sequences may be the only choice to disturb circRNA expression in *C. elegans*. Fortunately, CRISPR-Cas9 based genome editing is quite versatile and of high efficiency in *C. elegans* (Paix et al., 2015).

The circRNA-KO strains generated in this work did not show any obvious phenotypes in several assays, like locomotion, chemotaxis, lifespan, and aldicarb resistance (data not shown). Even if some phenotypes were observed, they should be carefully interpreted since the linear mRNA levels could also be changed (Figure 3D).

In *zip-2*, two short pairs of RCMs, 7 nt and 13 nt in length, were identified. To my best knowledge, they are the shortest identified *cis* elements that promote circRNA formation. Moreover, the 13-nt RCMs are highly conserved in the *zip-2* ortholog genes in five nematode species, suggesting their roles in promoting exon-skipping and back-splicing may be conserved.

Currently, two models have been proposed to explain the correlation between exon-skipping and circRNA formation (Jeck and Sharpless, 2014; Jeck et al., 2013): 1). RCM-promoted back-splicing produces circRNAs and y-shaped intermediates, which are further spliced to form skipped transcripts; 2). Exon-skipping produces skipped transcripts and lariat intermediates, which are further back-spliced to form circRNAs (Figure 6). For the first time, I show that RCMs are not only required for back-spicing but also promote exon-skipping. I further delineated that RCMs are not promoting exon-skipping through back-splicing, neither the other way. Instead, the two pathways are happening together, possibly competing with each other. I propose that RCMs in the introns not only bring the splice sites for back-splicing to proximity but also bring the sites for exon-skipping together, facilitating both processes simultaneously. Since the RCMs still exist in the intermediates of back-splicing and exon-skipping, they may function twice to promote further splicing/back-splicing in these intermediates (Figure 6).

Previous studies of RCMs’ roles in circRNA regulation or splice sites required for back-splicing were mainly based on plasmids in cultured cells. In this work, I show that *C. elegans* is a useful model for *in vivo* investigation of circRNA regulation.

## Acknowledgment

The NW1229 strain was provided by CGC, which is funded by NIH Office of Research Infrastructure Programs (P40 OD010440). I thank WormBase (https://wormbase.org/). I thank Mr. David Knupp and Dr. Pedro Miura from the University of Nevada, Reno, for instructions on northern blot. I thank Dr. Ichiro Maruyama and Dr. Pedro Miura for scientific discussions. I thank the members of Information Processing Biology Unit (Maruyama Unit) for their discussions and feedbacks. I thank DNA Sequencing Section of OIST for the help on RNA sequencing. I am grateful for the help and support provided by the Scientific Computing and Data Analysis section of Research Support Division at OIST. I thank Okinawa Institute of Science and Technology, Graduate School for financial support.

## Author Contributions

D.C. designed and conducted all the experiments, performed all the analysis, and wrote the paper.

## Materials and methods

### Worm maintenance

*C. elegans* Bristol N2 strain was used as the wild type. Worms were maintained using standard conditions on Nematode Growth Media (NGM) agar plates with *Escherichia coli* strain OP50 (Brenner, 1974) at 20°C or 25°C. Strains used in this study are listed in Table S1.

### Worm synchronization

Worm synchronization was performed by bleaching for large-scale worm preparation (L1 worms for dissociation). Briefly, worms were washed off plates using M9 buffer when a lot of eggs were laid and most of the worms were gravid adults. The worms were washed with M9 buffer and then bleached in bleach solution (1 M NaOH, 0.6% (m/v) NaClO) with ∼5 minutes continuous shaking. Then eggs were pelleted and washed three times with 12 ml M9 buffer by centrifuging at 2000 rpm for 0.5 minutes. Finally, the egg pellet was re-suspended in ∼5 ml M9 buffer and rocked at room temperature for 17-24 hours to hatch. For small-scale worm preparation (locomotion assay), worms were synchronized by egg-laying. Briefly, 10-15 gravid adult worms were placed onto NGM plate for four hours, and worms were removed after egg-laying. The eggs were then cultured to the desired stage.

### L1 worm dissociation

To ∼80 µl of L1 worm pellet, 200 µl SDS-DTT solution (200 mM DTT, 0.25% SDS, 20 mM HEPES pH 8.0, 3% sucrose) was added, followed by a 1.5-minute incubation at room temperature. Then, the worm pellet was washed 5x with 1 ml egg buffer (25 mM HEPES pH 7.3, 118 mM NaCl, 48 mM KCl, 2 mM CaCl2, 2 mM MgCl2, 0.340 ± 0.005 Osmolarity) and centrifuged at 10,000 ×g for 30 seconds. The washing steps and centrifugation should be performed quickly so that one round of washing and centrifugation is done in 40-50 seconds. The washed worm pellet was re-suspended in 100 µl pronase (15 mg/ml in egg buffer) from *Streptomyces griseus* (Sigma-Aldrich). Worms were dissociated by periodic mechanical disruption by pipetting for 15 minutes. 200 µl tips were used for mechanical disruption by the method mentioned in Zhang et al.’s protocol (“Pipette the larvae suspension with a 200 µl tip during the digestion. Adjust the pipetting volume to the approximate volume of the suspended pellet. Slowly pull suspended larvae into the pipette tip. Then, press down to force the pipette tip against the bottom of the microcentrifuge tube and slowly eject the contents”) (Zhang et al., 2011). Do as many times as possible. When most worm bodies were dissociated, 900 µL L-15/FBS medium (10% FBS in Leibovitz’s L-15 medium (Gibco), 0.340 ± 0.005 Osmolarity adjusted by sucrose) were added. Cells were collected and washed twice with 1 ml egg buffer by centrifuging at 9600× g for 5 min at 4°C. Cells were suspended in the appropriate amount of egg buffer and allowed to sit on ice for at least 30 min. The upper volume of cell suspension was used for FACS. For whole worm control, after dissociation and washing, the cell suspension was put on ice in the whole procedure of sorting.

### Fluorescence-Activated Cell Sorting (FACS)

Sorting was performed on a FACS AriaII flow cytometer (Becton Dickinson) equipped with a 70 µm nozzle. 2 µm and 3.4 µm polystyrene beads (Spherotech) were used for size calibration. Before sorting, propidium iodide (PI) was added to the cell suspension to a final concentration of 0.2 − 0.5 µg/ml. Then, profiles of dissociated cells from GFP-labeled strains were compared to profiles of cells from N2 worms to exclude auto-fluorescent cells. Sorted cells were collected in 3 ml L-15/FBS medium in a 15 ml conical tube chilled on ice. For RNA extraction, sorted cells and whole worm control samples were collected by centrifugation in a swing-bucket centrifuge at 4400 rpm, 4°C for 10 min. The supernatant was removed and 0.3 ml Trizol solution (Invitrogen) were added and stored at −80°C. For culture, sorted cells were seeded onto a poly-D-lysine coated glass-bottom dish (MatTec) with daily changes of L-15/FBS buffer. Cells were visualized by confocal microscopy (Carl Zeiss, LSM780) with a 60 × oil lens.

### Mutagenesis by CRISPR-Cas9

Mutation by CRISPR-Cas9 was based on the protocol published by Dokshin et al. (Dokshin et al., 2018) with minor modifications. Briefly, Cas9 (Sigma-Aldrich; 0.5 μl, 10 μg/μl in supplied buffer), tracrRNA (Sigma-Aldrich; 5 μl, 0.4 μg/μl in 10 mM Tris-HCl, pH7.5) and designed crRNA (ThermoFisher or IDT; 0.4 μg/μl in Tris-HCl, pH7.5; 2.8 μl for single crRNA, 1.4 μl each for 2 crRNAs) were mixed and incubated at 37°C for 10 min. Then recombinant dsDNA fragment (add to > 400 ng/μl final concentration) or recombinant single-strand DNA (ordered from Invitrogen or IDT; 2.2 μl, 1 μg/μl in Tris-HCl, pH7.5), injection marker pRF4(*rol-6*) (2.7 μl, 300 ng/μl in Tris-HCl, pH7.5), KCl (0.5 μl, 1 M), HEPES (1 μl, 0.2 M, pH7.4) and H2O were added to make a final 20μl injection mixture. Injected P0 worms were recovered at 20°C overnight and then transferred to RT (25°C). For F1 with obvious phenotypes, F1 worms with target phenotype were picked, and homozygous progeny were kept. For mutagenesis with no obvious genotypes, ∼10 F1 rollers were picked onto separated plates, and their genotypes were checked by single worm PCR after laying eggs. Large indels were identified by amplicon size differences. Small indels were checked by enzyme digestion of amplicons. Non-roller homozygous progenies with target genotype were kept. CrRNAs, recombinant ssODNs, validation primers, and restriction enzymes used in this work are listed in Table S3.

### RNA extraction

RNA extraction from sorted samples and whole worm samples was performed using Direct-zol RNA MicroPrep kit (ZYMO Research) with on-column DNase I (ZYMO Research) digestion according to the manufacturer’s protocol. RNA quality and quantity were measured by High Sensitivity RNA ScreenTape (Agilent) on TapeStation 4200 (Agilent). For RNA extraction from worms, worms were first flash frozen in Trizol solution (Invitrogen) in liquid N2 and then homogenized in by vortexing with glass beads (φ 0.1 mm) in Beads Cell Disrupter MS-100 (TOMY). If not mentioned, all the RNA samples used in this study were from L1 worms of indicated genotypes.

### RNA Sequencing

For RNA-seq of samples from sorted neurons (the sort group) and whole worms (the whole group), libraries were prepared using KAPA RNA HyperPrep kit with RiboErase (HMR) (KAPA biosystems) according to the manufacturer’s protocol. The RNA input was 150 ng and fragmentation conditions were 85°C for 5 min. Barcodes were introduced to each sample using KAPA duel-indexed adapters (KAPA biosystems). Length distribution of each library was determined by TapeStation 4200 (Agilent) using High Sensitivity DNA ScreenTape (Agilent). Libraries were quantified by KAPA library quantification kit (KAPA biosystems) and then multiplexed and sequenced on Illumina Hiseq 4000 platform to obtain 150 nt paired-end reads.

### Droplet digital PCR (ddPCR)

cDNA was reverse transcribed from 10 ng total RNA using an iScript Advanced cDNA synthesis kit (Bio-Rad). ddPCR was performed by using ddPCR EvaGreen Supermix kit (Bio-Rad) on a QX200 Droplet Reader (Bio-Rad) based on the manufacturer’s protocol. Results were analyzed using QuantaSoft software (Bio-Rad).

### Real-time PCR

Real-time PCR reactions were performed using soAdvanced Universal SYBR Green Supermix (Bio-Rad) with cDNAs synthesized from iScript Advanced cDNA synthesis kit (Bio-Rad). 20 μl reaction mix with 2 μl cDNA (∼1-10 ng) were monitored on StepOnePlus Thermal Cycler (Applied Biosystems) in “fast mode”. Cycling conditions: 95°C, 30’, 40 or 45 cycles of 95°C, 15’ and 60°C, 30’with plate reading, and a final melt curve stage using default conditions. Primers used are listed in Table S2.

### RNase R treatment

Total RNA was treated with or without (Mock) RNase R (2 U/μg) in the presence of Ribolock(2 U/μg) (ThermoFisher Scientific). The reaction was incubated at 37°C for 30 min. Then RNA was purified with an RNA Clean and Concentrator kit (ZYMO Research) according to the manufacturer’s protocol. For fold change quantification, RNA was quantified by Nanodrop and an equal amount of RNA input was used for cDNA synthesis. For northern blot, 20 μg total RNA with or without RNase R treatment was used for loading.

### Northern blot

Northern blot was performed using NorthernMax kit (ThermoFisher Scientific) and the probes were labeled by α-^32^P-deoxycytidine 5’-triphosphate (PerkinElmer) using Random Primer DNA Labeling Kit Ver. 2 (Takara, #6045) according to manufacturer’s protocols. Briefly, RNA samples (10 μg or 20 μg) were resolved in 1% agarose gel by electrophoresis at 5 V/cm in 1× MOPS buffer for ∼2 hours. Then RNA was transferred onto an Amersham Hybond-N+ membrane (GE Healthcare) by capillary blot for 2.5 hours using the transfer buffer supplied in NorthernMax kit. Transferred RNA was crosslinked by 254 nm UV at 1200 × 100 μJ/cm^2^(Analytik Jena CL-1000). Prehybridization was performed in ULTRAHybe buffer at 50°C for one hour, followed by hybridization with ^32^P labeled probes overnight at 50°C. The membrane was washed 2 × 5 min at room temperature using Low Stringency Washing Solution and 2 × 15 min at 50°C using High Stringency Washing Solution. The membrane was sealed in kitchen wrap and exposed to a phosphorscreen for several hours to overnight, and the signals were detected by Typhoon FLA7000 (GE Healthcare). Quantification of bands was performed using ImageQuant software (GE Healthcare). Primers used for probe amplification are listed in Table S2.

### circRNA prediction and RNA-seq data analysis

The DCC pipeline (Cheng et al., 2016) was used for the prediction of circRNAs with RNA-seq data. Briefly, raw reads were aligned to reference genome (WBcel235/ce11) using STAR (Dobin et al., 2013) (https://github.com/alexdobin/STAR) with the following options: -- outSJfilterOverhangMin 15 15 15 15 –alignSJoverhangMin 15 –alignSJDBoverhangMin 15 -- outFilterScoreMin 1 --outFilterMatchNmin 1 --outFilterMismatchNmax 2 --chimSegmentMin 15--chimScoreMin 15 --chimScoreSeparation 10 --chimJunctionOverhangMin 15. Then the output files from STAR, chimeric.out.junction, were used for circRNA annotation with DCC (https://github.com/dieterich-lab/DCC). Predicted circRNAs from DCC were filtered with at least three junction reads in each group. Differential expression analyses of mRNAs and circRNAs were performed using DESeq2 (Love et al., 2014) package in R with the gene count output from STAR or the BSJ junction count output from DCC, respectively. The plots (PCA plots, boxplots, scatter plots) were generated using ggplot2 package (https://ggplot2.tidyverse.org/), and ggpubr (http://www.sthda.com/english/rpkgs/ggpubr) package in R.

### RCM analysis

RCM analysis in flanking introns of circRNAs or non-circular control exons was performed using IntronPicker and autoBLAST scripts (https://github.com/alexandruioanvoda/) described in (Cortes-Lopez et al., 2018).

### Microscopy

Confocal images were obtained using a Zeiss LSM780 confocal microscope, and images were processed using the ZEISS ZEN3.1 software.

### Gene ontology analysis

Gene ontology enrichment analysis was performed using WormBase Enrichment Suite webserver (Angeles-Albores et al., 2018; Angeles-Albores et al., 2016) (https://wormbase.org/tools/enrichment/tea/tea.cgi).

## Statistical analysis

Statistical analysis was performed using R or Prism (GraphPad).

## Data availability

Raw FASTQ files from the RNA-seq data were deposited at the NCBI Sequence Read Archive (BioProject: PRJNA669379). All strains and other materials are available upon request.

## Supporting Figures

**Figure S1:**
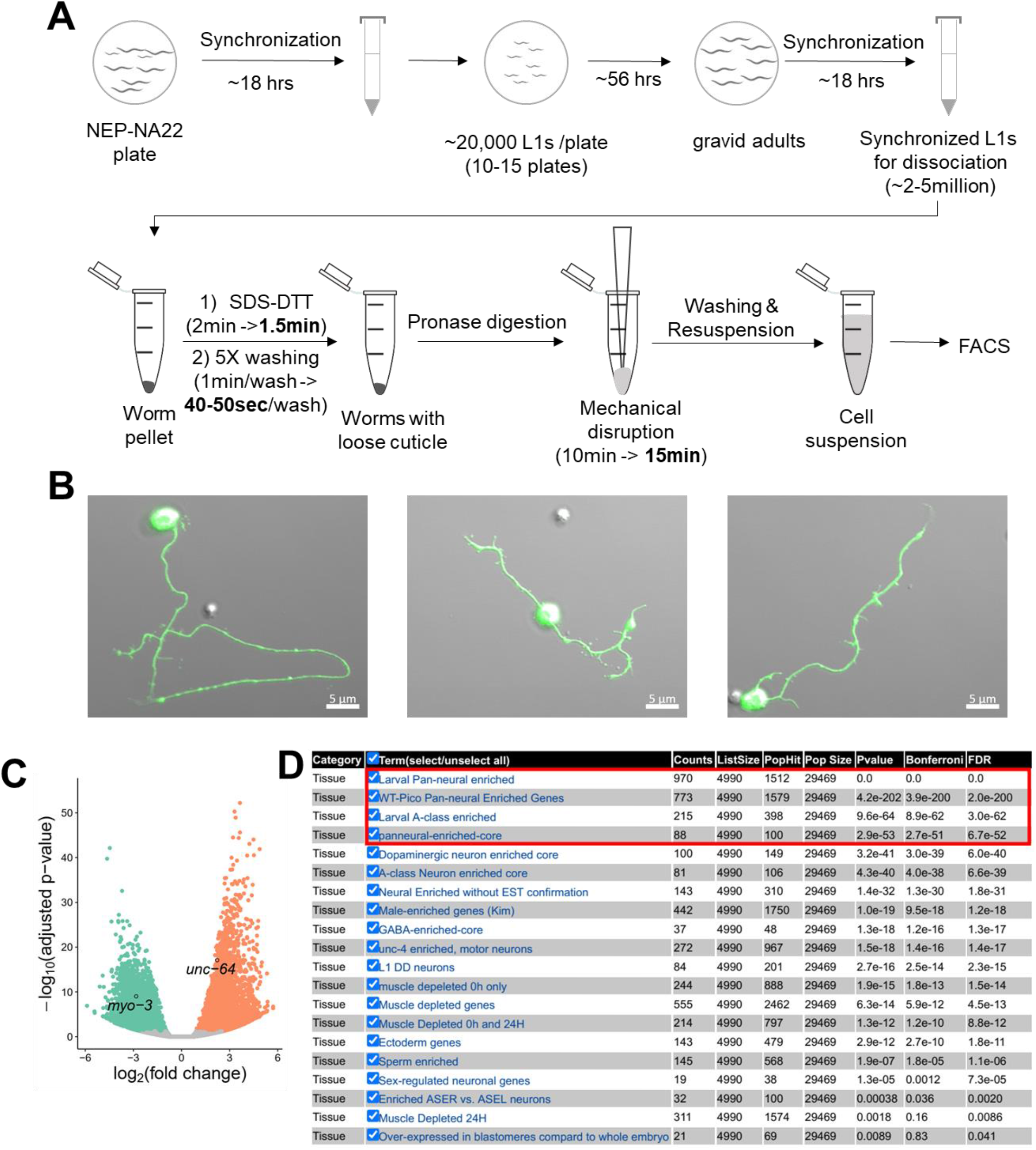
related to Figure 1. (A) Steps of L1 worm preparation and dissociation for FACS. Optimized conditions are in bold. (B) Confocal images showing sorted neurons after five-day culture at 20°C. Scale bars are 5 μm. (C) Volcano plot showing differentially expressed genes between the sort group and the whole group. *myo-3* and *unc-64* are labeled. (D) Output from WormExp for gene set enrichment search using upregulated genes in the neurons in our dataset. Red rectangle highlights the top 4 hits.

**Figure S2:**
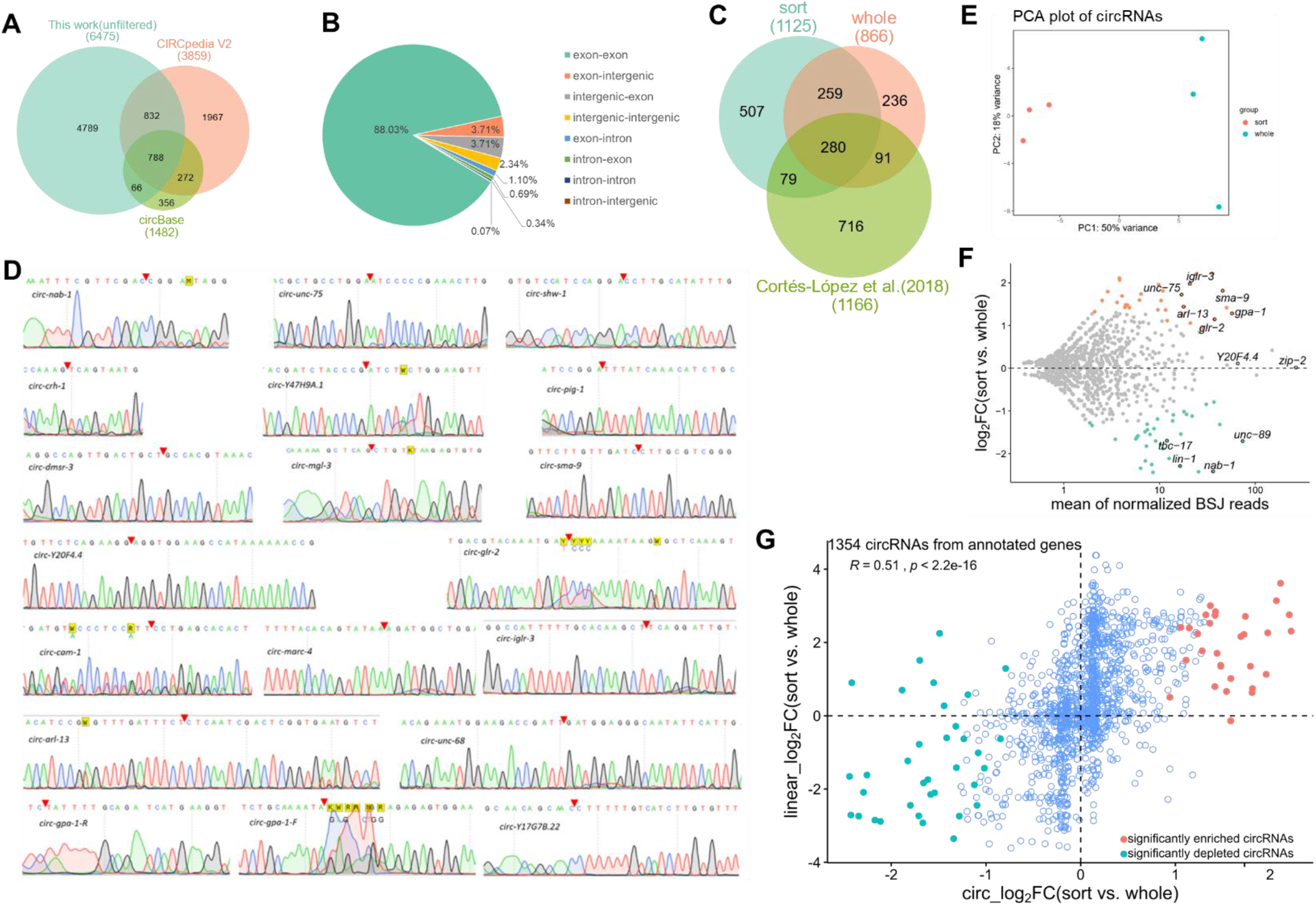
related to Figure 2. (A) Overlap of circRNAs identified in this work and circRNAs of *C. elegans* in two databases (CIRCpedia V2 and circBase). (B) Ratios of junction types of filtered circRNAs. (C) Overlap of filtered circRNAs in this work and filtered circRNAs in work of Cortés-López et al (*1*). (D) Sanger sequencing results of the BSJ sequences of selected circRNAs. Red triangles denote the joint sites. (E) PCA plot of circRNAs in the sort group and the whole group. (F) MA plot showing differentially expressed (DE) circRNAs between the sort group and the whole group. Significantly DE circRNAs are highlighted by colors. The gene names of some circRNA genes are labeled. (G) Scatter plot showing the correlation of log2 fold change of circRNAs and their cognate linear RNAs in the sort group and whole group. The Pearson correlation coefficient (*R*) and *p* value (*p*) are shown. Significantly DE circRNAs are shown by colored dots.

**Figure S3:**
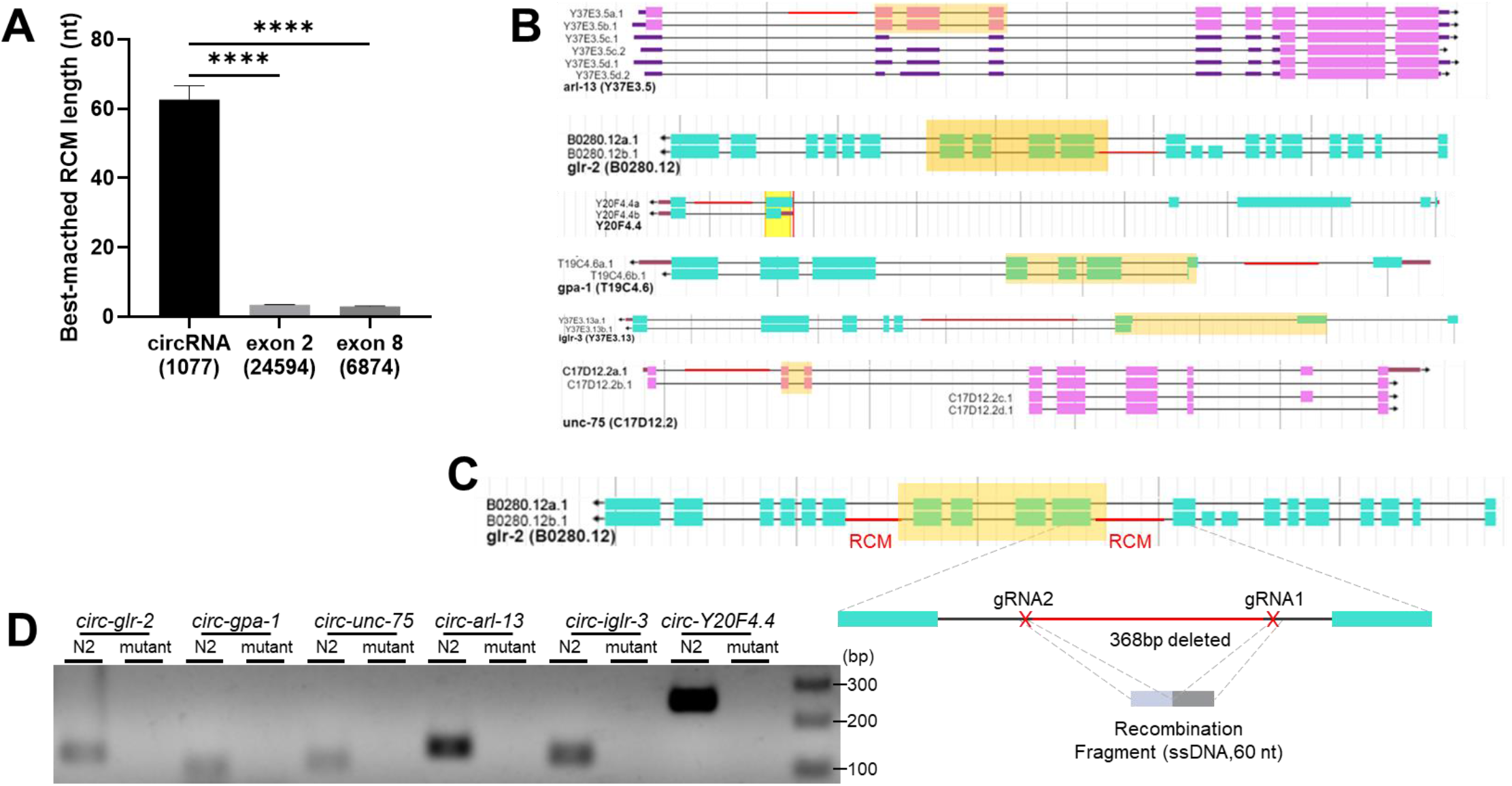
related to Figure 3. (A) Lengths of best-matched RCMs in one pair of circRNA-flanking introns compared with those in non-circRNA control exons (exon 2 and exon 8). Values are shown as mean ± SEM. Numbers in the brackets are numbers of intron pairs for analysis. *p* values are from Kruskal-Wallis test with Dunn’s post-hoc test for multiple comparisons. ****, *p* < 0.0001. (B) Positions of deleted RCMs (red line) of the 6 circRNA genes. Exons in orange shadows are to form circRNAs. (C) Illustration of RCM deletion in *glr-2*. Red lines in introns are RCMs. Red crosses denote gRNA positions. (D) Amplification of circRNA using divergent primers in wildtype N2 strain and RCM-deletion strains (mutant) of 6 circRNA genes.

**Figure S4:**
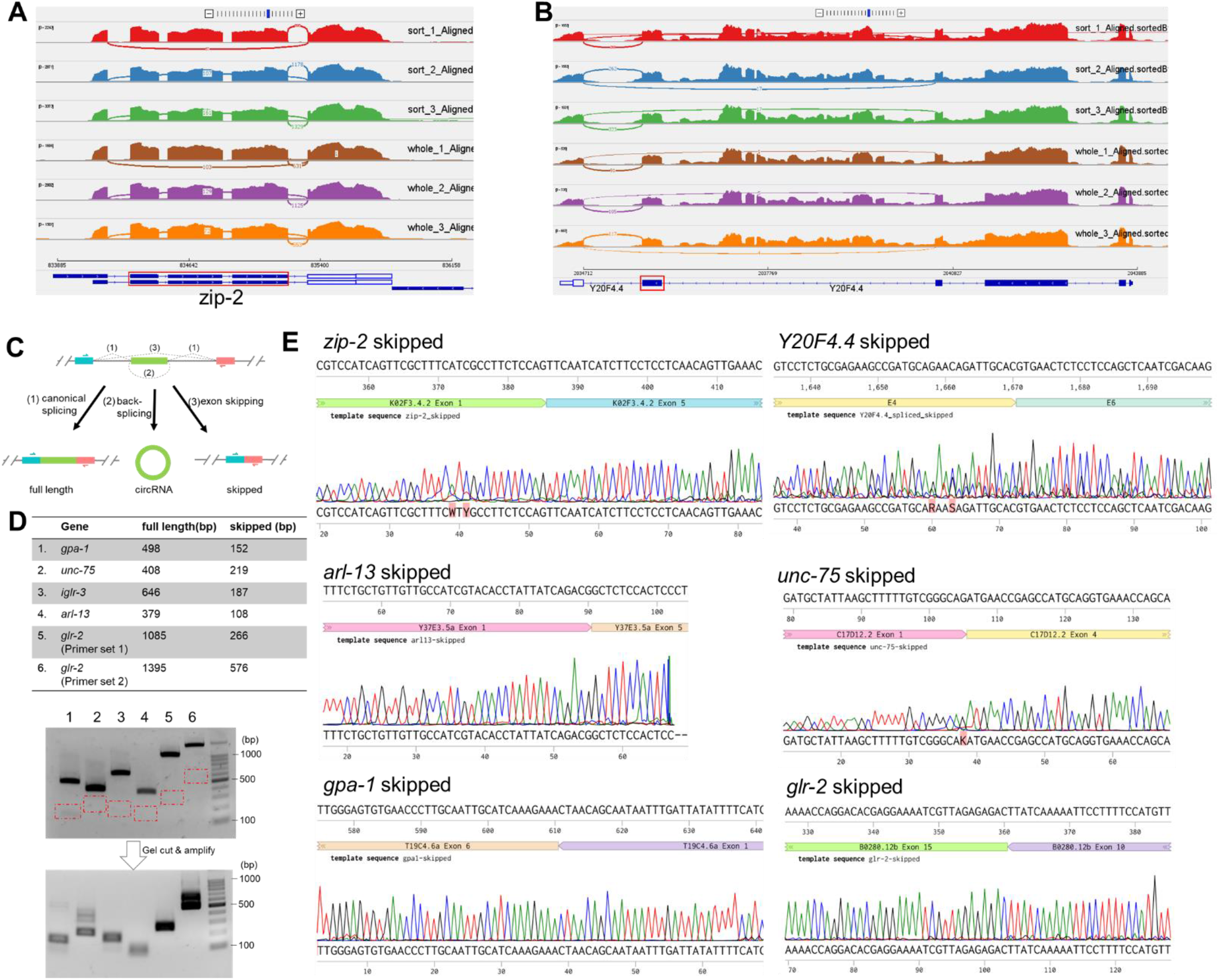
related to Figure 4. (A & B) Sashimi plots showing the number of exon-skipping junction reads in *zip-2* and *Y20F4*.*4* in our RNA-seq dataset. Exon(s) in red rectangles are circularized to form circRNAs. (C) Illustration of a circRNA-producing gene producing 3 transcripts: full-length, circular and skipped. Primers used to detect both the full length and the skipped transcripts are shown. (D) Amplification of the skipped transcripts from several circRNA genes by 2-round PCRs. Red rectangles mark the gel areas to be cut. (E) Confirmation of sequences of the skipped transcripts in 6 circRNA genes.

**Figure S5:**
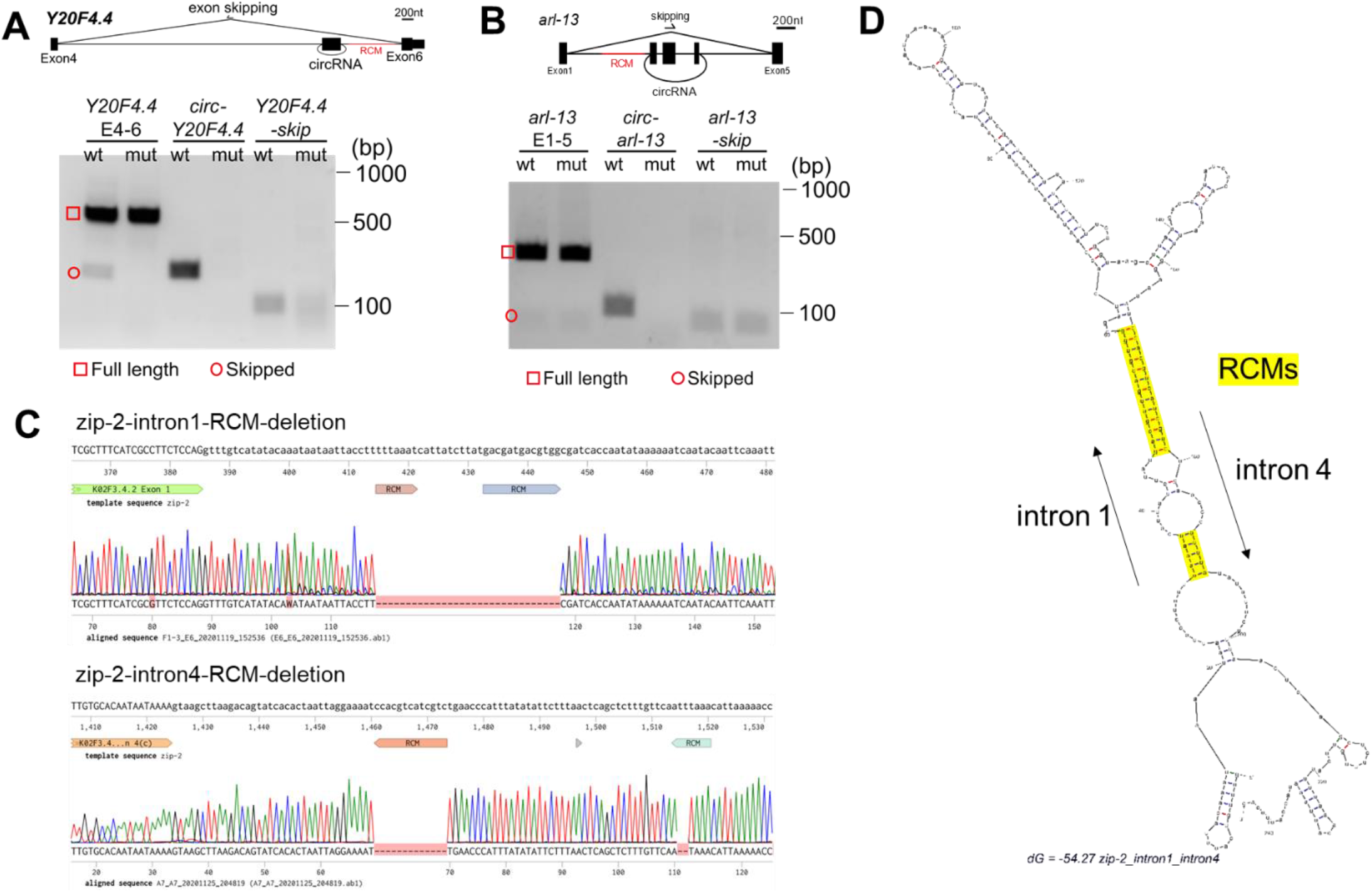
related to Figure 4. (A) RT-PCR detection of *Y20F4*.*4* transcripts in wildtype N2 strain (wt) and the RCM-deleted *Y20F4*.*4* strain (mut). (B) RT-PCR detection of *arl-13* transcripts in wildtype N2 strain (wt) and the RCM-deleted *arl-13* strain (mut). (C) Deleted RCM sequences in intron 1 and intron 4 of *zip-2*. (D) Folding prediction of intron 1 and intron 4 of *zip-2* by Mfold (http://www.unafold.org/mfold/applications/rna-folding-form.php). RCM sequences are highlighted.

**Figure S6:**
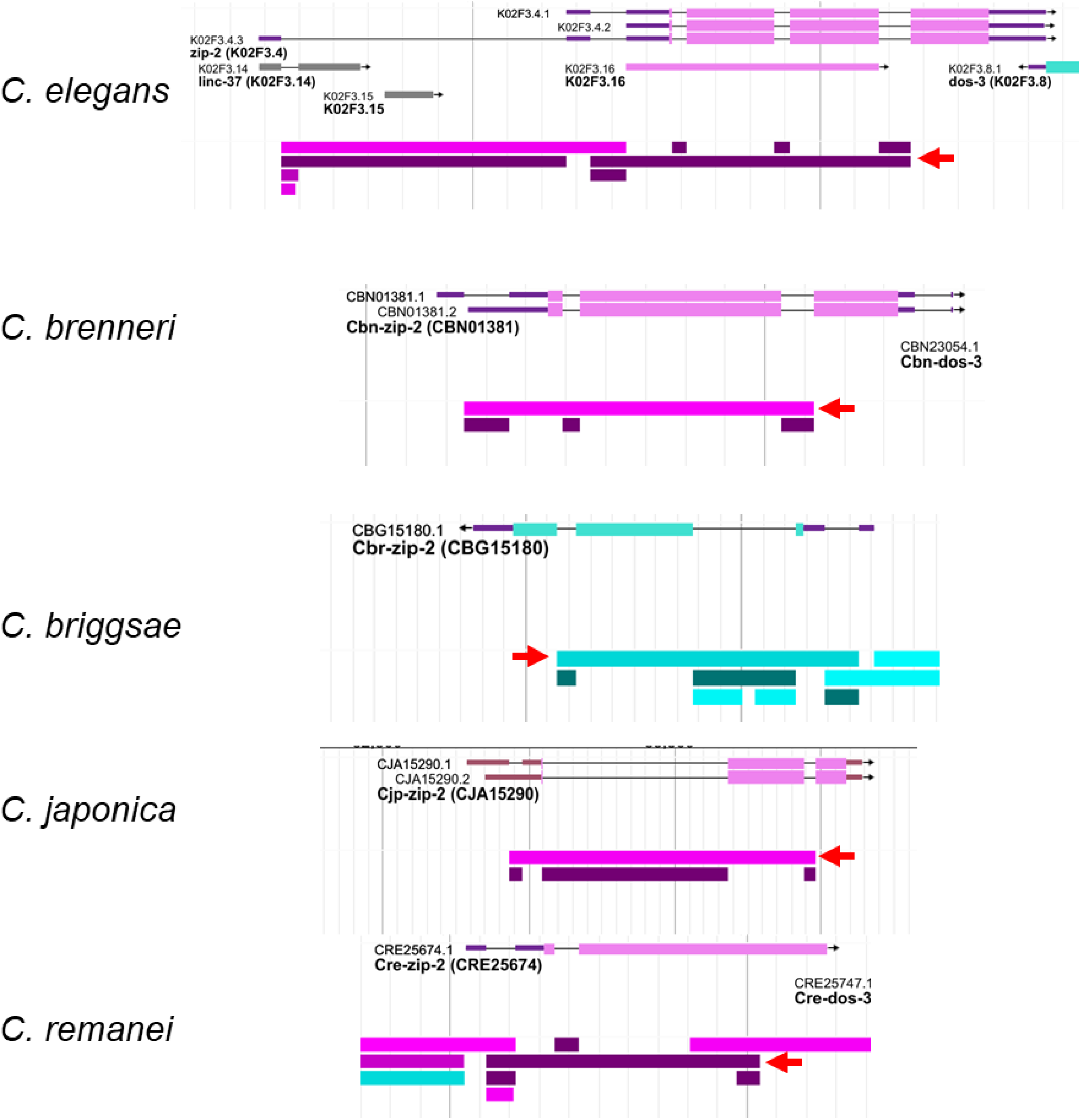
related to Figure 5. Gene structure of ortholog *zip-2* genes in indicated nematode species. The splicing patterns of these genes are also shown (from WormBase). Red arrows indicate the splice junctions of the skipped transcripts.

**Figure S7:**
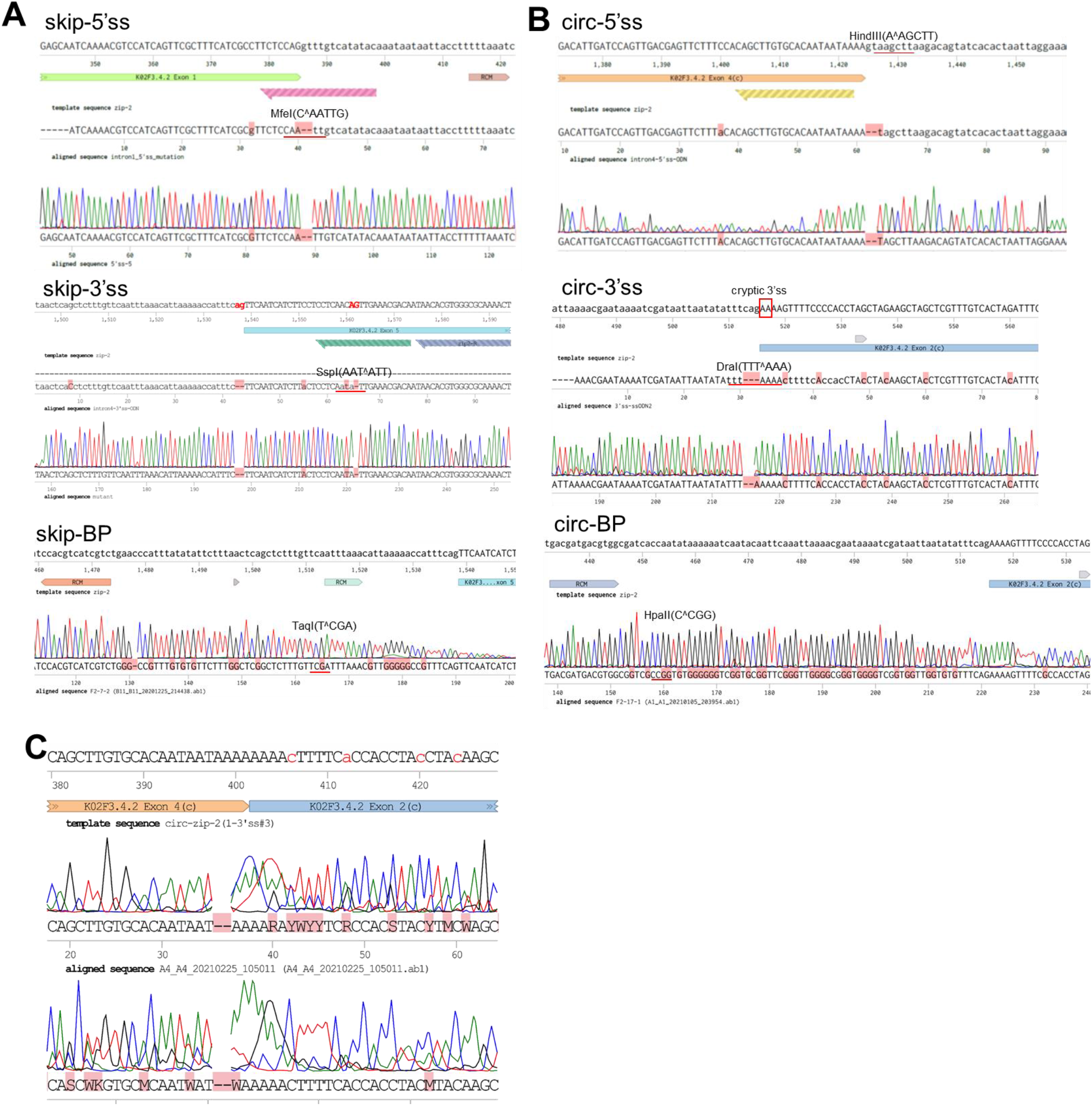
related to Figure 6. (A, B) Sanger sequence results of splicing sites and branch points mutation in intron 1 and intron 4 of *zip-2*. The enzyme digestion sites used to distinguish wildtype sequences and mutated sequences are labeled. Position of the cryptic 3’ss in circ-3’ss mutation is labeled. (C) Sanger sequence of circ-zip-2 produced from the zip-2(circ-3’ss) strain. Note that amplified sequences are 2 nt shorter than the predicted BSJ sequences.

